# Electron flow through NDH-1 complexes is the major driver of cyclic electron flow-dependent proton pumping in cyanobacteria

**DOI:** 10.1101/2020.09.21.307322

**Authors:** Neil T. Miller, Michael D. Vaughn, Robert L. Burnap

## Abstract

Cyclic electron flow (CEF) around Photosystem I is vital to balancing the photosynthetic energy budget of cyanobacteria and other photosynthetic organisms. The coupling of CEF to proton pumping has long been hypothesized to occur, providing proton motive force (PMF) for the synthesis of ATP with no net cost to [NADPH]. This is thought to occur largely through the activity of NDH-1 complexes, of which cyanobacteria have four with different activities. While a much work has been done to understand the steady-state PMF in both the light and dark, and fluorescent probes have been developed to observe these fluxes *in vivo*, little has been done to understand the kinetics of these fluxes, particularly with regard to NDH-1 complexes. To monitor the kinetics of proton pumping in *Synechocystis* sp. PCC 6803, the pH sensitive dye Acridine Orange was used alongside a suite of inhibitors in order to observe light-dependent proton pumping. The assay was demonstrated to measure photosynthetically driven proton pumping and used to measure the rates of proton pumping unimpeded by dark ΔpH. Here, the cyanobacterial NDH-1 complexes are shown to pump a sizable portion of proton flux when CEF-driven and LEF-driven proton pumping rates are observed and compared in mutants lacking some or all NDH-1 complexes. It is also demonstrated that PSII and LEF are responsible for the bulk of light induced proton pumping, though CEF and NDH-1 are capable of generating ∼40% of the proton pumping rate when LEF is inactivated.

**Highlights statement:** NDH-1 is essential for proton pumping during cyclic photosynthetic electron flow in cyanobacteria

## 1. Introduction

NADH dehydrogenase complex I, or complex I, is a widely distributed bioenergetic complex, with homologs seen in archaea, bacteria, and eukaryotes [1]. The core structure of these complexes and their subunit composition have been largely conserved throughout these phyletic kingdoms, with the bacterial complexes possessing the fewest subunits, and eukaryotes possessing accessory subunits, but retaining a core that is conserved across the domains of life [2-7]. These complexes are vital parts of a diverse variety of electron transport chains and have been shown to couple proton pumping to electron transport from ferredoxin (Fd) or NAD(P)H to a quinone to increase the proton motive force (PMF) for ATP production in heterotrophic bacteria as well as mitochondria and chloroplasts [3, 8-13]. These complexes have also been shown to act in reverse to dissipate PMF while producing low potential reductant (e.g. NAD(P)H), though this has only been observed under special inhibitor conditions with the addition of succinate in mitochondria and submitochondrial particles [14] or in chemoautotrophic bacteria and archaea with unique energy budgets such as *Nitrospira* or *Thaumarchaea* respectively [1, 15-18].

Cyanobacteria also possess complex I, however they are unique in possessing four functional versions of complex I, and they are termed NDH-1 for <u>N</u>ADPH <u>D</u>e<u>h</u>ydrogenase <u>1</u> complex, though they have recently been shown to utilize Fd for their reductant over NADPH [7, 19, 20]. Recently, the structure of two cyanobacterial NDH-1 complexes have been solved [4-7], covering representatives of the major functions of these complexes: <u>C</u>yclic <u>e</u>lectron flow (CEF), respiratory electron flow, and CO_2_ uptake. A summary of the components of photosynthetic electron flow, their actions on proton pumping, and inhibitors utilized here are shown in **Fig. 1A**. These NDH-1 complexes share a common “core” subcomplex, NDH-1M, with modular attachments that confer a specific function, such as CO_2_ uptake [4-6, 20, 21] (**Fig. 1B**). The core subcomplex contains the redox active subunits that participate in electron transfer from Fd^red^ to the <u>p</u>lasto<u>q</u>uinone (PQ) pool, and it pairs with specific NdhD and NdhF proteins in the membrane arm to assemble the functional complexes [4-6, 20] (**Fig. 1B**). These assembled complexes participate in CEF around <u>P</u>hoto<u>s</u>ystem I (PSI) by accepting electrons from Fd^red^ produced through photosynthetic electron flow and passing them into the PQ pool [22, 23] while presumably pumping protons to contribute to PMF, though proton pumping by the cyanobacterial NDH-1 complexes had not been experimentally demonstrated until this work. The NDH-1 complexes contribute majorly to CEF in cyanobacteria, but it has also been shown that another major route is through succinate dehydrogenase (SDH) [24], though its CEF activity is able to be limited based on the illumination, inhibitor additions, and mutations in the strains measured [25]. The PQ pool electrons are then passed through the photosynthetic electron transport chain, including cytochrome B_6_f Q-cycles, to pass them back to PSI and inevitably to Fd^ox^/NADP^+^, thereby keeping the redox state of [Fd^red^/NADPH] relatively constant while, it is hypothesized, contributing to PMF to drive [ATP] synthesis. Quantitative information on the thylakoid PMF in cyanobacteria has been obtained using electron spin resonance probes [26] and by measuring the distribution of radioactive pH-sensitive probes [27, 28]. These have provided a relatively consistent picture with the cyanobacterial cytoplasm at approximately pH 7.0 and a lumen pH of about 5.0-5.5 in the dark, supported by respiration. Photosynthetic electron transport was observed to increase the pH in the cytoplasm and decrease the pH in the lumen by approximately 0.5 pH units each, thereby increasing the ΔpH by about one unit [26]. This would produce a lower lumen pH than the more moderate values estimated for chloroplast thylakoids [29], nevertheless is in line with responses seen in plants and algae. Importantly, the electrical and proton concentration components of the PMF, ΔΨ and ΔpH respectively, were shown to be interconvertible using classical techniques employing ionophores and protonophores [28]. The steady-state pH of the lumen in the light appears to be regulated by various mechanisms allowing for photosynthetic electron transport while preventing over-acidification and allowing for photosynthesis to continue with minimal ROS production (reviewed in [30, 31]).

**Figure 1.**
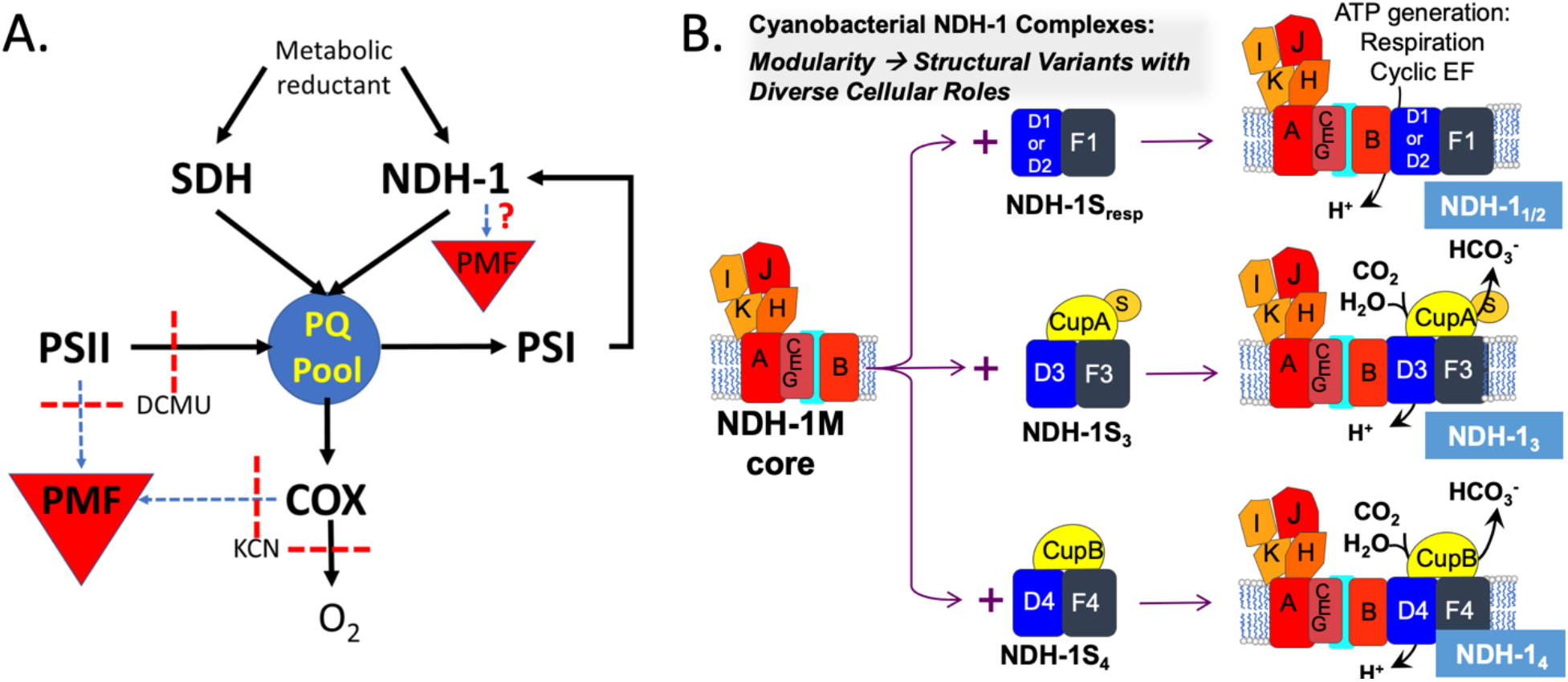
Photosynthetic electron flow and proton pumping in cyanobacteria. **Panel A**. The enzymatic components are shown, connected by arrows representing electron transfer. Blue dashed arrows indicate contribution to PMF, and red dashed lines represent a blocking of electron flow or proton pumping upon addition of an inhibitor. **Panel B**. The modularity of the cyanobacterial NDH-1 complexes. They share a common core (NDH-1M) containing NdhB,A,B,C,E,G,H,I,J,K, but differ in their distal membrane subunits, and are named based on the proteins they possess. The variable moduelsd provide two major functions that are largely split between the NDH-1_1/2_ and NDH-1_3/4_ complexes, which are primarily involved in CEF/respiration and CO_2_ uptake, respectively. The CEF/respiration complexes (NDH-S_resp_) modules contain the NdhD1 (or NdhD2), NdhF1 proteins plus smaller polypeptides, whereas the CO_2_ uptake modules contain either NdhD3/NdhF3/CupA/CupS for high affinity CO_2_ uptake or NdhD4/NdhF4/CupB/ for low affinity CO_2_ uptake.

The four cyanobacterial NDH-1 complexes are termed here based on the identity of the NdhD protein they possess, as each complex has an unique NdhD protein associated with it. Of the four complexes in cyanobacteria, two of them, the NDH-1_1_ and NDH-1_2_ complexes, are primarily active in CEF and respiratory electron flow [22, 24]. While the NDH-1_1/2_ complexes share a common NdhF protein termed NdhF1, the physiological differences between the NDH-1_1/2_ complexes are largely unknown. It has been shown to that the deletion of *ndhF1* severely inhibits CEF, and that NDH-1_1/2_ complexes are responsible for the bulk of NDH-1 driven electron flow [22]. These complexes have also been shown to form super complexes with PSI in cyanobacteria as a means of protecting against stresses such as high light or salt or in the absence of PSII activity after treatment with DCMU, a PSII inhibitor [32, 33]. It was shown that the NDH-1-PSI supercomplex stabilizes PSI [33], with the function of the supercomplex likely being to consume electrons for CEF as quickly as possible, limiting the space needed for Fd^red^ to diffuse [32]. This accelerated consumption of electrons is thought to act as an antioxidant mechanism in this way, especially when stresses such as high light leads to increased Fd reduction. Recently, it was shown that exposure to high light caused heavy accumulation of the NDH-1_3/4_ complexes, indicating their CO_2_ uptake activity in addition to their CEF activity may play a role in the NDH-1 antioxidant mechanism [7]. The deletion of the NDH-1_1/2_ complexes causes a hyper-oxidation of the PQ pool, locking the photosynthetic machinery in State 1 where the bulk of light harvesting is focused around PSII [34, 35]. Reduction of the PQ pool can trigger a “state transition” to State 2, where the bulk of light harvesting is focused around PSI. These state transitions are dependent upon the redox state of the PQ pool, and, when the photosynthetic electron transport chain is intact, there is normally a state transition upon the dark-light shift from State 2 to State 1 [36-38]. This suggests that the NDH-1_1/2_ complexes are important for mediating the reduction of the PQ pool both in the dark during respiration and in the light during photosynthesis. The tightly regulated response of cyanobacteria to these stresses highlights the importance of this mechanism for the protection and acceleration of PSI-CEF when needed, thereby providing electron transport pathways necessary for photosynthetic metabolism.

The other two complexes, NDH-1_3_ and NDH-1_4_, possess unique subunits that have carbonic anhydrase activity, termed <u>C</u>O_2_ <u>U</u>ptake or “Cup” proteins, and they participate in the <u>C</u>O_2_ <u>C</u>oncentrating <u>M</u>echanism (CCM), which functions to increase the internal concentration of inorganic carbon (C_i_) to 1000-fold the external concentration, hydrating CO_2_ to HCO_3_^-^, thereby trapping it in the cytoplasm [39, 40]. These two complexes differ in their gene expression, affinity for CO_2_, and flux of CO_2_ hydration, with the NDH-1_3_ complex being essential for growth in air [41]. They contribute relatively little to CEF compared to their NDH-1_1/2_ cousins, but do still participate [22]. These complexes require all of their unique components in order for CO_2_ uptake to occur, and lacking them or a component of the core subcomplex, NdhB, results in an inability to grow in limiting concentrations of CO_2_ [21, 41].

Evidence of proton pumping in cyanobacterial NDH-1 complexes and their physiological contributions to that activity will provide relevant data in terms of accounting for photosynthetic electron flow driven proton pumping, as well as provide insight into the mechanism of the unique cyanobacterial CO_2_ trapping system of the NDH-1_3/4_ complexes. While the e^-^ transport activity of NDH-1 complexes in general has been well probed [22-25, 42, 43], many questions remain in understanding the coupling of this electron transport to their potential proton pumping activity. Do NDH-1 complexes pump protons? Is that activity light dependent? Given the multiplicity of forms of cyanobacterial NDH-1 complexes, which are responsible for the bulk of proton pumping activity? Does CEF couple to proton pumping? Here, a pH sensitive fluorescent dye called <u>A</u>cridine <u>O</u>range (AO) that locates to both the cytoplasm and thylakoid lumen [44] was utilized to observe the formation of ΔpH across the thylakoid membrane in strains deficient in NDH-1 activity. This was done alongside chlorophyll (Chl) fluorescence measurements for the comparison of proton pumping activity with photosynthetic electron transport activities. In this work, it is hypothesized that CEF drives proton pumping, respiratory-like NDH-1_1/2_ complexes are responsible for the bulk of CEF driven proton pumping, and that CEF is incapable of reaching the rate of proton pumping reached by LEF.

## 2. Methods

### 2.1 Strains and molecular constructs

Strains of *Synechocystis* sp. PCC 6803 (hereafter, *Synechocystis*) were maintained on pH 8 BG-11 [45] with 1.5% agar supplemented with 18mM sodium thiosulfate and containing the appropriate antibiotics for each mutant (7µg/mL Gentamycin (Gm) for Δ*ndhF1*, 50µg/mL Kanamycin (Km) for M55), with WT maintained under ∼70 µE m^−2^ sec^-1^ light, Δ*ndhF1* 70 µE m^−2^ sec^-1^ light, and M55 50µE m^−2^ sec^-1^ light. Experimental material was obtained from 100mL cultures grown in 250mL Erlenmeyer flasks with rotary shaking (200 rpm) under ∼100µE m^−2^ sec^-1^ Cool White fluorescent lighting (GE). Cultures were harvested upon reaching an OD_750_ 0.7-1.0. The M55 strain was grown under 5% CO_2_ with an autoclaved gassing apparatus consisting of a 0.2µm filter attached to tubing put through a foam cap, sterilized, used to replace the foam cap of the regular shaking BG-11 cultures, with a long enough tube to gently bubble the media with CO_2_ while shaking.

The plasmid used to construct Δ*ndhF1*, pUCF1-Gm, was produced using Gibson assembly [46] by amplifying ∼500bp upstream and downstream regions of the *ndhF1* gene from wild-type chromosomal DNA utilizing the primers shown in **Table S1**. The Gm antibiotic resistance cassette was amplified from *Synechocystis* HT-3 genomic DNA [47] with the primers also shown in **Table S1**. The primers above were designed with overlaps for Gibson Assembly according to manufacturer’ s specifications (NEB). The three PCR generated fragments were assembled in the presence of pUC18 digested with NdeI and PstI (NEB) by Gibson Assembly according to the manufacturer specifications to create pUCΔF1. M55 was provided to our group by professor Teruo Ogawa from Nagoya University in Nagoya, Japan.

### 2.2 Chlorophyll Fluorescence

Cells were harvested by centrifugation, washed with 50mM Tricine pH 8 + 25mM NaCl (TCN) and resuspended to 5.9µg/mL Chl, measured by in a UV-Vis spectrophotometer (Shimadzu) [48]. Following this, each sample was diluted to 5µg/mL Chl in the cuvette with the Tricine buffer. Measurements of Chl fluorescence yield were measured using the Dual PAM-100 (Walz) with the 9-AA/NADPH module. For 5-minute illumination period measurements, a nominal light intensity of 53µE m^−2^ sec^-1^ was utilized except in experiments measuring LEF saturation as a function of light intensity, or experiments observing fluorescence in saturating light. Values for q_p_ were produced using the equation outlined in [49] utilizing multiple turnover flashes (300 milliseconds, nominally 20,000 µE m^−2^ sec^-1^).

### 2.3 Acridine Orange Fluorescence Quenching

Acridine orange (AO) has been used as a fluorescent dye for measuring proton pumping across a membrane in both whole cyanobacterial cells and membrane vesicles [44, 50-52]. Cells were grown and washed with TCN buffer as above for Chl fluorescence. Samples were diluted to 5.9µg/mL Chl had acridine orange added to 5µM, and were incubated shaking gently in the dark for 20 minutes to allow acridine orange penetration into the cell [44]. Samples of the cells were then added to a cuvette, and 1M KCl added to 150mM to dilute the cells to 5µg/mL and provide K^+^ to exchange across the thylakoid when Valinomycin (Val) is added to 10µM to dissipate ΔΨ. DCCD was added to 500µM to inhibit ATP synthase and prevent proton leakage. KCN was added to 200µM to inhibit cytochrome oxidase (COX) and respiratory proton pumping as well as to maintain the photosynthetic electron transport chain in a State 2 like poise [34]. Samples were then stirred for 5 minutes in the dark, and the fluorescence measured with the JTS-100 (Biologic) or Dual-PAM 100 (Walz). A 534/20nm bandpass filter (Edmund Optics) was placed between the sample and the detector in both instruments.The 9-AA/NADPH attachment (Walz) was used on the Dual-PAM 100 to excite AO, however, the wavelength used does not allow for maximal excitation [53], unlike with the JTS-100.

The Dual-PAM 100 proved useful for measuring dark fluorescence and some larger scale changes in AO fluorescence, while the JTS-100 was primarily used to observe rates of proton pumping in response to illumination. Dual-PAM 100 data was generated by exporting the data averaging 100 points, resulting in one point per 100ms, with inhibitors (Val, DCCD, KCN) added in the dark. Upon reaching a steady-state, a new measurement was performed on the same sample with a 5 minute actinic illumination with multiple turnover flashes (parameters above) before, during, and after illumination. The multiple turnover flashes were included for measurement of Chl fluorescence parameters at the same time (**Fig. S1**), and were not greatly taken into account for the AO measurements.

For the AO fluorescence data recorded on the JTS-100, points were collected once per second in the dark and once per 100ms in the light, with light intensity varying by experiment. Each sample was recorded after a 5 minute dark incubation after the addition of inhibitors with 5 minutes of additional stirring in the dark between each technical replicate. The rates and depths of quenching were relatively unaffected by repeated measurements or a slight drift in these conditions (discussed below). The inhibitors Val (10μM), KCN (200μM), and DCCD (500μM)were added to samples measured in the JTS-100 to ensure the measurement of maximal rates, as discussed below. Accordingly, these formed the standard set of inhibitors for most experiments used to observe proton pumping here. Utilizing the same conditions in the instrument, DCMU was added in some case to the sample to 10µM and measurements recorded after a 5 minute incubation period in the dark with stirring, and measurements recorded as described above. A fresh sample was then prepared with the standard inhibitors along with the addition of 10µM DCMU and 250µM CCCP, incubated for 5 minutes with stirring, and measurements recorded as described above. To obtain sufficient signal/noise ratios, 3 to 6 technical replicates were recorded per condition for each of 3 biological replicates. AO data collected on the JTS-100 was provided by averaging the data of all 3 biological replicates for each condition with a line plotted through the data after about 4 seconds of illumination and extending for 4-10 seconds, depending on the shape of the trace and the ability to plot a line through the linear portion of quenching. The traces were arbitrarily zeroed to the beginning of the linear portion of quenching. Rates were determined by plotting the line through each biological replicate and using those to determine the average rate and standard deviation.

## 3. Results

### 3.1 Proton pumping activity in whole Synechocystis sp. PCC6803 cells

In cyanobacteria, the pH sensitive dye AO has been shown to report the changes in pH across the thylakoid membrane by fluorescence changes [44, 52]. The cyanobacterial thylakoid membrane presents a unique challenge because both photosynthesis and respiration occur in the same membrane and utilize the common electron carriers [54, 55]. Therefore, a suite of inhibitors was used to isolate the contributions of specific complexes, including NDH-1 complexes, to ΔpH. Firstly, a significant fraction of PMF is stored as a K^+^ ion gradient, contributing to the electrical component, ΔΨ, of PMF [26, 28, 52], and there is a considerable efflux of K^+^ ions upon illumination, to create a positive ion deficit in the thylakoid lumen to drive the accumulation of H^+^ [56]. The ΔΨ component can cause variations in proton pumping measurements using AO [52], therefore, Val was used to dissipate the K^+^ gradients across the membranes with the addition of 150mM KCl to remove the ΔΨ component of PMF from measurements. ATP synthase was inhibited with 500µM DCCD to prevent the flow of protons through the membrane from affecting the measurements. Finally, in some cases, 200µM KCN was added to block COX, thereby pushing dark-adapted cells into a State 2-like poise, where more light harvesting occurs at PSI than in State 1, since the NDH-1 mutants used have the potential to be locked in State 1 [34]. The addition of KCN also allows for the isolation of the photosynthetic components of ΔpH formation as there is a considerable the ΔpH in the dark when COX is active [57]. The activity of COX in the dark results in a certain amount of ΔpH [57] (**Fig. 2A-C**).

**Figure 2.**
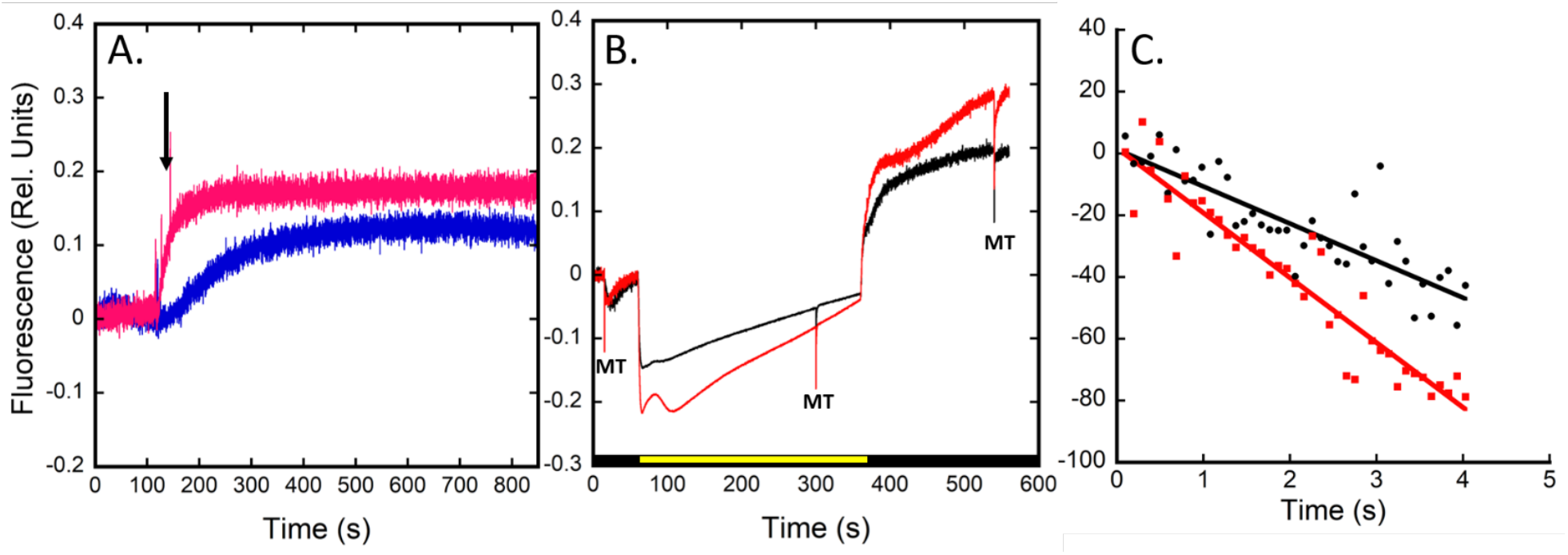
Respiratory electron flow contributes to ΔpH in the dark. **Panel A:** Acridine fluorescence in the Walz DUAL PAM-100 instrument. Cells were dark adapted with acridine orange (5µM) in TCN buffer with KCl (150mM). Either only KCN (200µM) (blue trace) or KCN plus Val (10µM) and DCCD (500µM) (pink trace) were added (arrow). **Panel B:** Acridine fluorescence in the DUAL PAM-100. Cells were incubated in the dark for 5 min. with background inhibitors in the absence (black) or presence (red) of KCN and AO fluorescence was collected over the course of a 5 min. illumination period, including multiple turnover flashes before, during, and after illumination. The MT flashes were employed for evaluating characteristics of the simultaneously measured Chl fluorescence (e.g. Fig. S1). Data was acquired using same instrument settings and scale, and the traces zeroed to F_0_ to account for small drifts in baseline and sample differences. **Panel C:** Acridine fluorescence in the BioLogic JTS-100 instrument. Cells were dark adapted with acridine orange (5µM) for 20 min, then a sample prepared with KCl (150mM) and either Val (10µM) and DCCD (500µM) (black circles), or with the inclusion of KCN (200µM) (red squares) and the sample stirred in the dark for 5 min. Traces are typical of three replicates.

The presence of the dark ΔpH provides PMF which must be overcome in order for photosynthetically driven proton pumping to occur. The electrochemical back-pressure of positive charges within the thylakoid lumen ultimately has an effect on the ability of light induced proton pumping [52], so dissipating these pressures can allow observation of photosynthetically driven proton pumping unimpeded by the effects of that back-pressure. The contribution of respiratory activity in dark-adapted cells to their trans-thylakoid PMF can be observed as an increase in AO fluorescence (loss of quenching), as the dark ΔpH dissipates upon the addition of KCN as detected using a PAM fluorometer due to the loss of COX activity (**Fig. 2A**). This dissipation is enhanced when Val and DCCD are included with KCN. The addition of any of these alone did not result in as strong a dissipation as when all three are included (not shown), indicating the combination of these additions is necessary for the disruption of dark ΔpH. These results are indicative of the dissipation of dark PMF associated with respiration. This is further shown in **Fig. 2B**, where AO fluorescence arbitrarily zeroed to the fluorescence before illumination is seen in the absence and presence of KCN in addition to the background inhibitors. In the presence of KCN, quenching occurs to ∼1.5x the value seen in its absence, and occurs about twice the rate (**Fig. 2C**). This suggests that respiration in the dark indeed exerts some amount of electrochemical back-pressure that impedes the rate ΔpH formation upon illumination. Slower kinetic processes are also apparently occurring simultaneously. Superimposed upon the rapid light-induced AO quenching, we observe a more gradual increase in AO fluorescence (**Fig. 2B**). This can be attributed to the previously observed alkalinization of the cytoplasm occurring in the minutes time frame, potentially due to CBB-cycle activity [26-28, 58].

Comparison of Chl fluorescence induction kinetics with and without KCN revealed a reduction of the PQ pool in the dark, and some alterations in the rates of secondary transitions, likely involving S-state transitions and the activation of the CBB cycle, but otherwise photosynthetic electron transport is unaffected, save for the dramatic rise in both F_0_ and post-illumination fluorescence upon addition of KCN, likely due to the lack of COX as an electron sink exacerbating the actinic effect of the PAM measuring light and thwarting the consumption of photosynthetically-produced reductant during the light-to-dark transition (**Fig. S1**).

### 3.2 Proton pumping activity dominated by photosynthetic linear electron flow

To evaluate the activity of different photosynthetic membrane complexes with respect to proton pumping, initial rates of light-induced AO fluorescence quenching in the presence of the three background inhibitors (KCN, Val, DCCD), were tested in a shorter experiment. Steady state AO fluorescence in the dark, F_0_, was recorded for 20 sec, followed by quenching due to the application of 600 µE m^−2^ sec^-1^ actinic illumination for 15 seconds, followed by darkness to observe the recovery phase (**Fig. 3**). The quenching of fluorescence upon illumination indicates the acidification of the thylakoid lumen due to the activity of photosynthetic electron flow upon a dark-to-light transition. The rate of this quenching during the linear phase after illumination(∼4-8 or 4-14 sec depending on the strain and additional inhibitors added) is termed *j*_H_^+^. The quenching occurred from F_0_ to an extent, termed F_ΔpH_ (**Fig. 3**). Cessation of illumination led to an increase in AO fluorescence indicating a collapse of ΔpH across the thylakoid [44]. This rise appears to be biphasic, occurring first rapidly and then slowly. There is additionally some upward drift in fluorescence seen after the cessation of illumination. As this is not seen in dark adapted cells (**Fig. 2A**), it is hypothesized that this may be due to the processing of reducing equivalents produced during illumination. Upon addition of DCMU to WT cells, the rate of acidification decreases to ∼40% that of the control (**Table 1**), however a considerable gradient is still formed, albeit to smaller extent in the time measured (**Fig. 3**). The inclusion of the uncoupler CCCP with DCMU causes a cessation of light-induced proton pumping, giving a baseline against which to compare the other data. DCMU was always added with CCCP; since, unless PSII is inhibited, the solution slowly acidifies due to water oxidation activity (not shown). These data indicate that LEF, including the strong contribution of protons released from water oxidation, is the major contributor to proton pumping during the dark-to-light transition, although the contribution of CEF is very substantial, at 40% of LEF.

**Table 1.** Relative rates of acidification. The rate of acidification, *j*_H_^+^, upon illumination in WT, Δ*ndhF1*, and M55 in the presence of the background inhibitors with either no additional inhibitors, with DCMU, or DCMU and CCCP added. Values are averages of three biological replicates with 3 averaged technical replicates each. Data is presented with the standard deviation. Data normalized to [Chl]/OD_750_ of the WT to account for differences in the Chl content per cell in the different strains.

| Inhibitors added | Wild-type | $\Delta ndhF1$ | M55 |
| --- | --- | --- | --- |
| Val+DCCD+KCN | $33.77 \pm 8.09 \text{ s}^{-1}$ | $23.24 \pm 6.99 \text{ s}^{-1}$ | $20.25 \pm 4.45 \text{ s}^{-1}$ |
| Val+DCCD+KCN+DCMU | $14.07 \pm 2.33 \text{ s}^{-1}$ | $0.72 \pm 0.59 \text{ s}^{-1}$ | $-0.70 \pm 0.47 \text{ s}^{-1}$ |
| Val+DCCD+KCN+DCMU+CCCP | $0.46 \pm 0.22 \text{ s}^{-1}$ | $-0.99 \pm 0.07 \text{ s}^{-1}$ | $-1.46 \pm 1.16 \text{ s}^{-1}$ |

**Figure 3.**
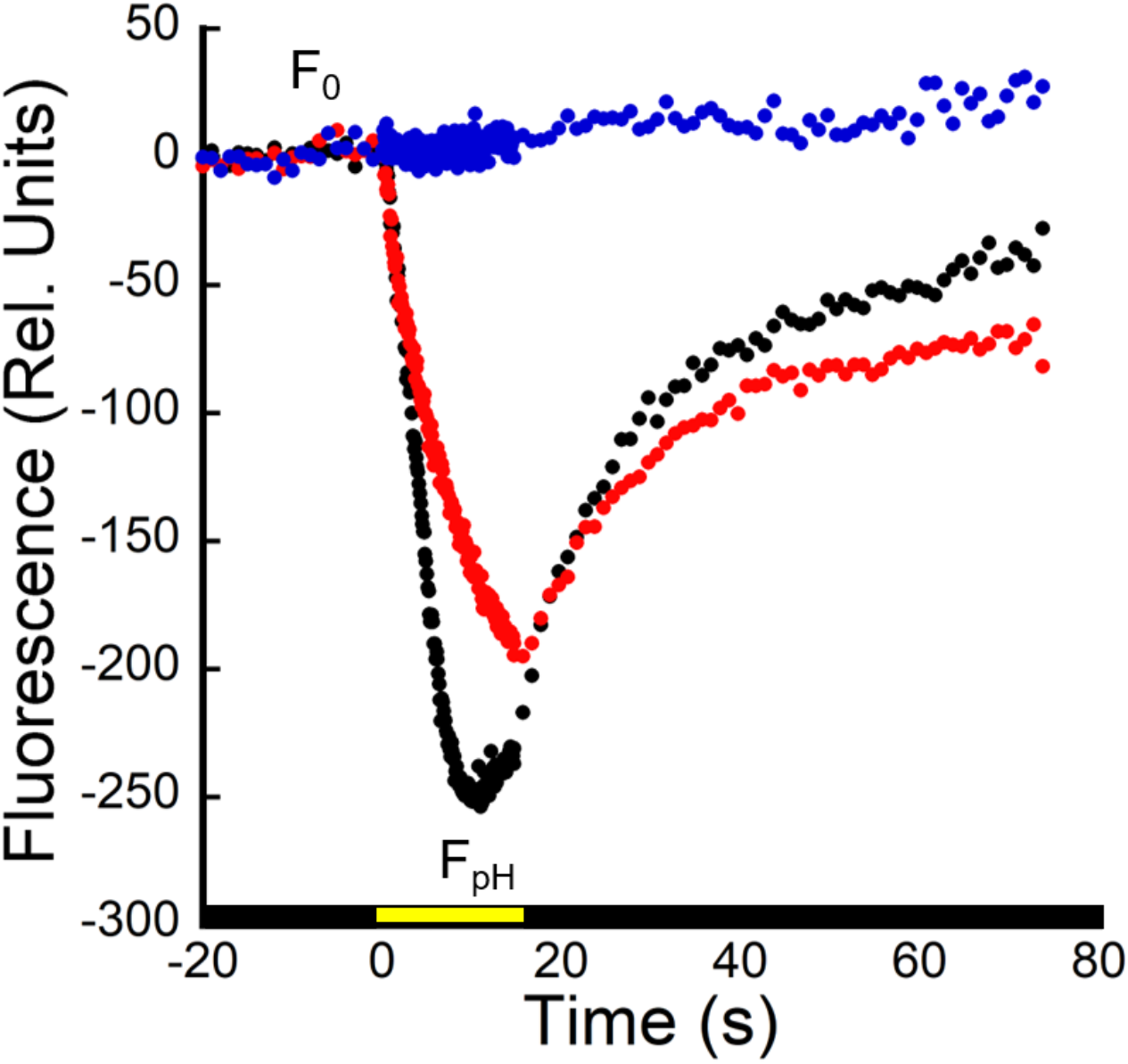
Light-induced acidification of the thylakoid lumen. Acridine orange fluorescence of dark-adapted WT cells in the JTS-100 representing the formation and dissipation of ΔpH across the thylakoid lumen upon the activation and cessation of actinic illumination. Cells were dark adapted with acridine orange (5µM) for 20 min, then a sample prepared with KCl (150mM) plus Val (10µM), DCCD (500µM), and KCN (200µM) and stirred in the dark for 5 min. Actinic illumination (630 nm, 600 μE) was applied for 15 seconds. Black symbols, standard additions; red symbols, standard additions plus DCMU, blue symbols, standard additions plus DCMU and the uncoupler CCCP. The plots are typical of three biological replicates, each with three technical replicates. Steady state fluorescence in the dark is labeled F_0_. Steady-state fluorescence in the light is termed F_ΔpH_. The linear portion of the section of fluorescence quenching between F_0_ and F_ΔpH_ is used to calculate the rate of quenching (*j*_H_^+^).

### 3.3 Proton pumping rate is dependent upon light intensity

To examine the components of proton pumping activity in the dark-to-light transition, the WT was measured in *j*_H_^+^ as a function of light intensity to observe the dependence of proton pumping on photosynthetic performance. As can be seen in in **Fig. 4A, C**, increasing light intensity caused an increase in the depth of AO quenching reached within the time measured. The rates and depth of quenching only increased to a point, and intensities greater than 300µE m^−2^ sec^-1^ did not appreciably increase the depth of quenching. Upon addition of DCMU to the cells (**Fig. 4 B, D**), blocking PSII water oxidation, there is no appreciable difference seen between the different light intensities in the depth of quenching. The rate of proton pumping in the WT when DCMU is added is ∼40% of the rate when PSII is active (**Table 1**), highlighting the importance of water oxidation in the formation of ΔpH in the light. Because the quenching is light dependent, it is indicative of photosynthetic activity, however this activity is apparently at its maximum even at lower light intensities. When plotting the rates of the linear portions of AO quenching, the clear dependence of the rate of proton pumping on light intensity can be observed (**Fig. 5A**). As reflected in the depth of quenching without DCMU, the rate of quenching does not appreciably increase at intensities greater than 300μE, indicating saturation of the rate of light-induced proton pumping. This is reflected by the proportion of open PSIIs, q_p_, which shows that PSIIs are largely occupied when ≥300μE light was applied (**Fig. 5B, Fig. S2**). When DCMU was added and the rates plotted (**Fig. 5A**, red squares), very little difference is seen between the rates regardless of light intensity. At even 70µE m^−2^ sec^-1^, CEF dependent proton pumping is at approximately maximal value, with minimal variation seen as light intensities climb, and none significantly different from one another. This indicates that CEF driven proton pumping, while dependent upon light to operate, is incapable of compensating for the lack of PSII. To determine whether this inability to compensate was due to stoichiometric limitations of CEF machinery or a lack of substrates to participate in electron transport, glucose was added to DCMU treated cells. Previous work has shown that addition of glucose to cells significantly increases the amount of reductant available for P700 re-reduction [59]. Interestingly, the addition of glucose had very little effect on DCMU treated cells, in fact slowing the proton pumping rate down mildly (**Fig. S4**). This indicates that even without glucose CEF-driven proton pumping is saturated at low light intensities, and that the difference in predicted versus observed proton pumping activity of CEF to LEF is due to the available complexes and their catalytic parameters, not a lack of reductant. While these results indicate that CEF indeed drives proton pumping, lending further credence to a long-standing hypothesis, further experiments are needed to determine which complexes contribute to CEF-driven ΔpH formation in the dark-to-light transition, as well as their relative contributions.

**Figure 4.**
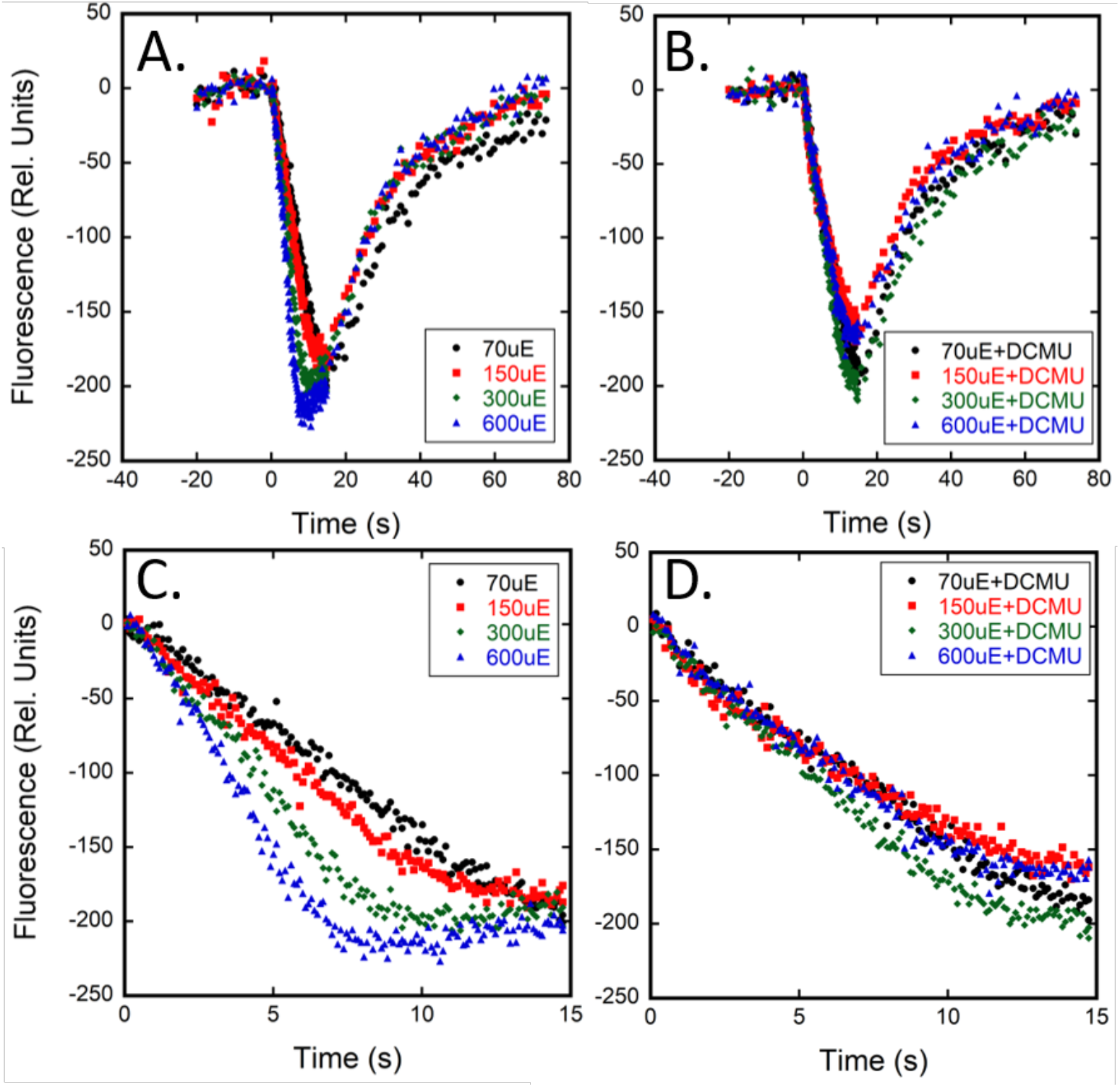
Light dependent proton pumping in the absence (A,C) or presence (B,D) of DCMU at varying light intensities. Acridine orange fluorescence of dark adapted WT cells in the JTS-100 in response to increasing light intensities, representing the formation and dissipation of ΔpH across the thylakoid membrane in response to the dark-to-light and light-to-dark transitions respectively. Cells were dark adapted with acridine orange (5µM) for 20 min, then a sample prepared with KCl (150mM) plus Val (10µM), DCCD (500µM), and KCN (200µM) and stirred in the dark for 5 min. For panels C and D, 10µM DCMU was used. Actinic illumination (630 nm, 600 μE) was applied for 15 seconds. These plots are typical of 3 technical replicates. **Panel A**: The formation and dissipation of light dependent ΔpH formation when LEF and CEF are both active. **Panel B**: The formation and dissipation of light dependent ΔpH formation when only CEF is active. **Panel C**: A closer look at the light dependent formation of ΔpH when LEF and CEF are active. **Panel D**: A closer look at the light dependent formation of ΔpH when only CEF is active.

**Figure 5.**
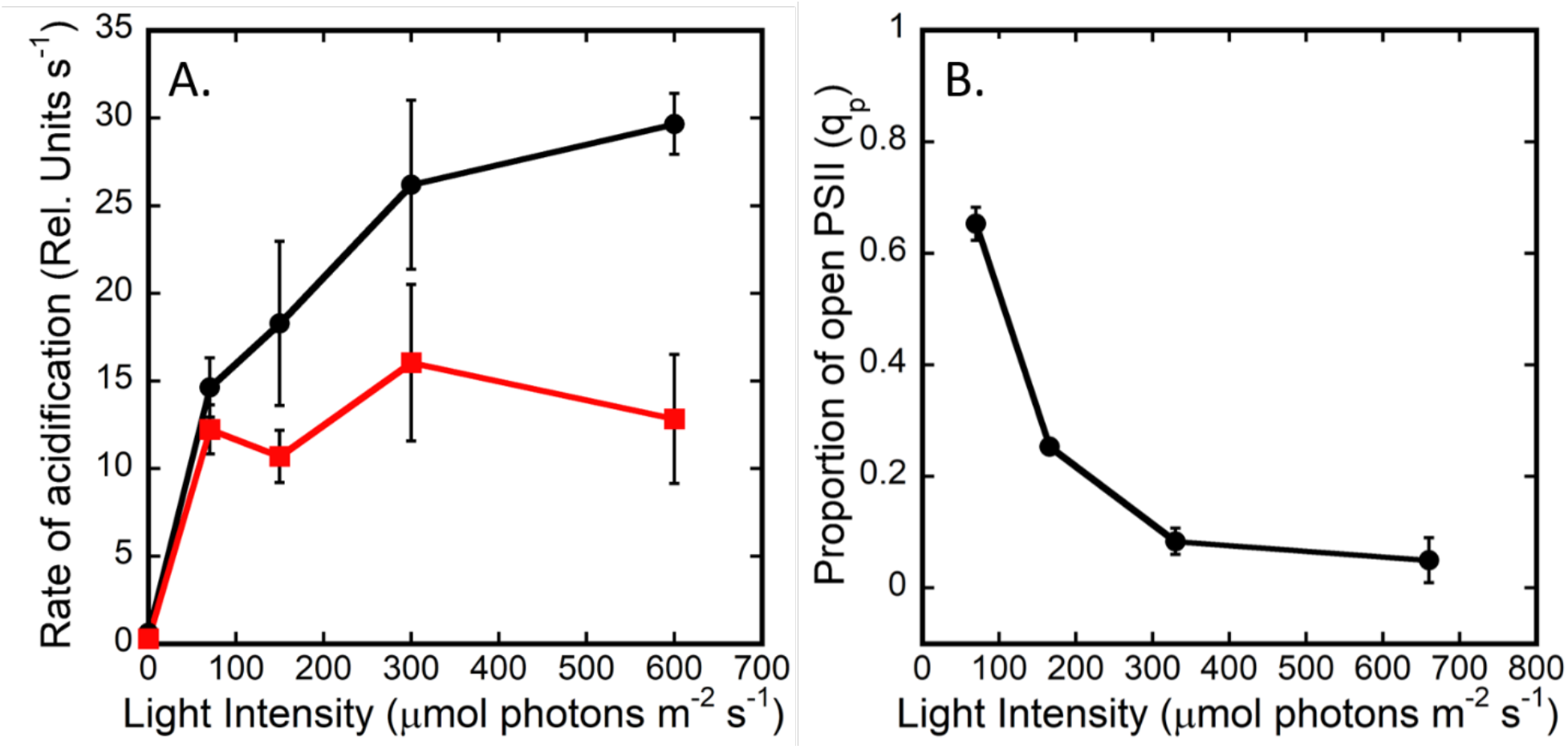
Proton pumping rates and photochemical quenching (q_p_) as a function of light intensity. **Panel A:** Rates of proton pumping as a function of light intensity determined from JTS-100 data in the WT without (black circles) or with 10µM DCMU (red squares). WT cells were dark adapted in TCN buffer with AO (5µM), KCl (150mM), Val (10µM), DCCD (500µM), and KCN (200µM). Actinic illumination (630 nm, 600 μE) was applied for 15 seconds. Each point is representative of the rate estimates of 3 biological replicates with 3 technical replicates. **Panel B:** The proportion of open PSII reaction centers as a function of light intensity as estimated as photochemical quenching (q_p_), in cells suspended to 5µg/mL Chl in TCN buffer + 150mM KCl. Photochemical quenching (q_p_=(F_m_-F_s_)/F_m_-F_0_)) was obtained from Chl fluorescence yields using a PAM fluorometer [49].

### 3.4 NDH-1_1/2_ complexes are largely responsible for NDH-1 driven proton pumping

To determine which NDH-1 complexes are responsible for CEF-driven proton pumping, mutants in the cyanobacterial NDH-1 complexes were utilized in proton pumping experiments. The mutants Δ*ndhF1* and M55, lacking functional NDH-1_1/2_ and all NDH-1 complexes respectively, were utilized to observe whether these complexes pump protons, and what contribution they have to proton pumping in the dark-to-light transition. As can be seen in **Table 1** and **Fig. 6**, there is little difference between the strains when PSII is active, further highlighting its importance in the establishment of ΔpH upon illumination, however, there is a steady decline in the rates as NDH-1 complexes are removed, with M55 having the slowest rate. The rates of proton pumping in WT and M55 with the background inhibitors are significantly different (*p*=0.03), indicating the NDH-1 complexes may be important for proton pumping in the dark-light transition, even when PSII is active. Upon the inhibition of PSII, the WT shows a slowing of *j*_H_^+^ to about 40%, however the formation of ΔpH in the Δ*ndhF1* and M55 strains is dramatically slowed (**Table 1, Fig. 6**). While Δ*ndhF1* is still able to form a gradient at ∼5% the rate of its no DCMU control, the M55 strain remains approximately in line with conditions measured with the protonophore CCCP near baseline (**Fig. 6B**,**C**, red and blue symbols respectively). When Chl fluorescence of the WT and Δ*ndhF1* was measured, it was seen that, in accordance with previous work [34], chlorophyll fluorescence in the dark increases in the presence of KCN, indicating a more reduced PQ pool (**Fig. S5**). M55, on the other hand, was relatively unaffected by the addition of KCN. As well, the characteristic post-illumination fluorescence rise associated with NDH-1 activity [7, 23, 41, 60] was observed in both the WT and Δ*ndhF1*, with its magnitude being enhanced by the addition of KCN, though to a much greater degree in the WT both with and without KCN. M55 had no rise in either condition. Together these results indicate that in the dark-to-light transition, the NDH-1_1/2_ complexes are major contributors to CEF-driven proton pumping in addition to their established role in electron flow, and they may indicate that the NDH-1_3/4_ complexes are also involved, but to a lesser extent.

**Figure 6.**
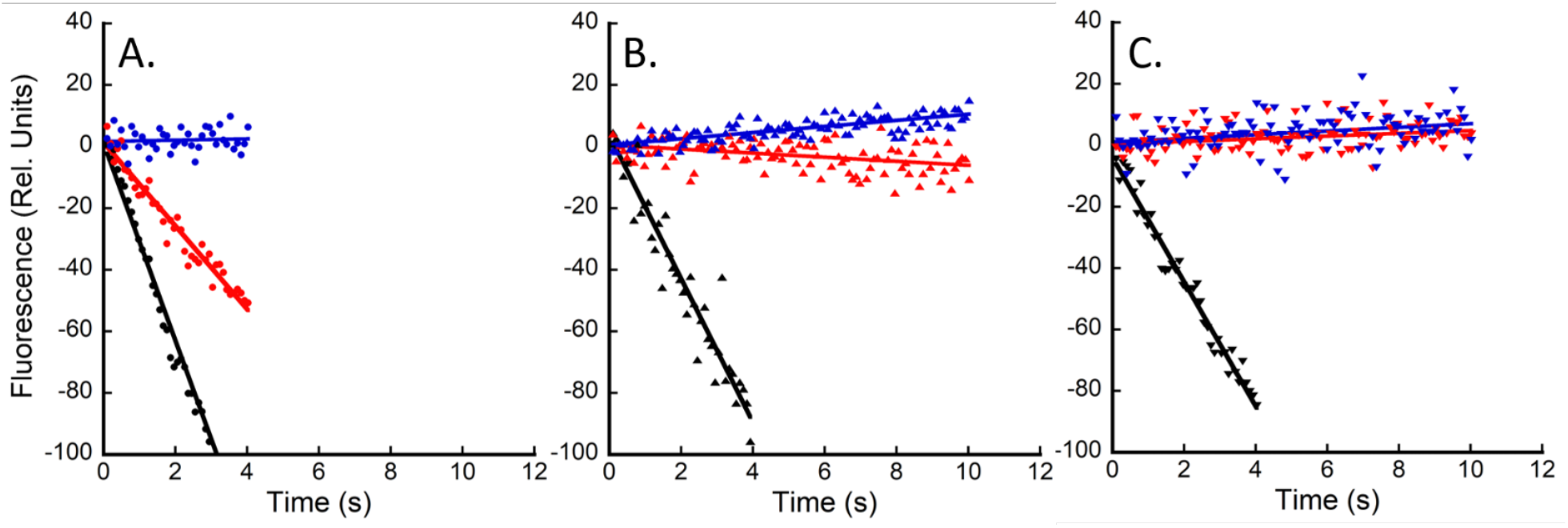
Rates of acidification, *j*_H_^+^, upon illumination in the WT and mutants deficient in NDH-1 complexes. Acridine orange fluorescence in the JTS-100. Cells were dark adapted in TCN buffer with acridine orange (5µM) for 20 min, then a sample prepared with KCl (150mM) plus Val (10µM), DCCD (500µM), and KCN (200µM) and stirred in the dark for 5 min. Actinic illumination (630 nm, 600 μE) was applied for 15 seconds. These plots are typical of 3 technical replicates. WT (**A**), Δ*ndhF1* (**B**), and Δ*ndhB* (**C**) after the addition of background inhibitors (black symbols), with the addition of 10µM DCMU (red symbols), and with the addition of DCMU and 250µM CCCP (blue symbols). Data is representative of the averages of 3 biological replicates with at least 3 technical replicates each. Data was normalized to [Chl]/OD_750_ of the WT to account for differences in [Chl] per cell.

## 4. Discussion

Since the discovery that PMF and ion gradients are central to virtually all bioenergetic processes [61], there has been considerable work done to understand certain characteristics of the cyanobacterial PMF both in the thylakoid and the cytoplasm [26-28]. Despite this work, the kinetics of proton pumping in cyanobacteria have been little explored with regards to the contributing protein complexes and the impacts of those components on rates of gradient formation. Here, the analysis of proton pumping by *Synechocystis* has revealed components that contribute to the formation of ΔpH upon the illumination of the cells by actinic light. An interesting advantage that whole cell cyanobacteria have over heterotrophic species in this sort of measurement is that the photosynthetic electron transport chain is light inducible, allowing for the control of redox components by controlling the actinic illumination. This allows for the relatively easy assaying of photosynthetic ΔpH formation in whole cell cyanobacteria, which provides opportunities to observe the effects of cellular metabolism on the proton gradient.

To best observe the contribution of the photosynthetic complexes to light-dependent proton pumping in whole cells, various parameters needed to be considered, especially as the focus of these experiments is to compare rates of photosynthetically driven proton pumping, particulary in regard to the NDH-1 complexes. It has been shown that the amount of K^+^ available in the thylakoid has effects on the ability of the cells to perform proton pumping, providing an electrochemical back-pressure against which protons must be pumped as they both have a positive charge [52]. Therefore, to alleviate this constraint, Val was added to the samples. To prevent the forward or reverse action of ATP-synthase from deducting from or contributing to the formation of the gradient, the inhibitor DCCD was also included. Because respiration occurs in the dark in cyanobacteria, it is reasonable to suppose that the dark ΔpH will also affect the observed rate and amplitude of light-induced AO quenching since it will have start from a ‘pre-energized’ membrane exerting an electrochemical backpressure as photosynthetic electron flow initiates. The existence of the dark ΔpH was demonstrated when cells were treated with KCN, causing the dark ΔpH to collapse, an effect enhanced by the addition of Val and DCCD alongside KCN (**Fig. 2A**). Indeed, by eliminating the dark ΔpH the rate and depth of light-induced AO quenching increased (**Fig. 2B**,**C**), indicating that in the absence of a a ‘pre-energized’ membrane, proton pumping is faster and that it is light-dependent proton pumping. This supports the conclusion that light-induced proton pumping can be limited, under in vivo conditions, to some degree by the electrochemical back-pressure due the dark ΔpH. Although AO fluorescence technique is not amenable to pH calibration, previous estimates of the dark ΔpH in cyanobacteria in the presence of Val is approximately 1.5 pH units and that value further increased to ∼2.5 pH units under illumination [26-28]. The observation that actinic illumination of dark-adapted samples in the presence of KCN produced a larger and more rapid quenching of AO fluorescence than in its absence (**Fig. 2B**) is consistent with these estimates. Overall, removing the dark ΔpH and ΔΨ allows for the determination of rates of light-dependent proton pumping that are closer to maximal within live cells.

While observing the major components of photosynthetically driven PMF formation, PSII was seen to be the major contributor to ΔpH formation in all strains measured. This is not an unexpected result, as the water oxidation reaction produces 4H^+^/O_2_, and PSII is both abundant in the cell and has a fast reaction rate compared to NDH-1 of other organisms [8, 21, 62-64]. This is demonstrated here by the sharp reduction in the rates of ΔpH formation upon the addition of DCMU in all strains tested (**Fig. 3, Fig. 4, Fig. 5A, Fig. 6, Table 1**), with that reduction most pronounced in strains lacking NDH-1 complexes. While the rates of proton pumping when PSII is active are similar in all the strains, there is a small reduction in the rates as NDH-1 complexes are removed. Nevertheless, the rates remain fast in all the strains, indicating that the NDH-1 complexes are important for proton pumping even when LEF is active, though their contributions are partially masked by LEF and PSII activity, as discussed below. The removal of PSII activity by the treatment of the WT with DCMU still allows for a substantial gradient to be formed upon illumination, indicating that CEF drives the formation of ΔpH in the absence of PSII activity, though less effectively as the rate drops to ∼40% (**Table 1, Fig. 3, Fig. 5A, Fig. 6A**). While the rate is decreased, it quenches to approximately 80% the F_ΔpH_ without DCMU, potentially indicating that the reduced rate is incapable of completely overcoming the dissipation mechanisms that allow F_ΔpH_ to be established (**Fig. S3**). AO quenching in DCMU treated cells is shown to be light dependent (**Fig. 3, Fig. 4, Fig.5A**), indicating that it is due to CEF activity.

The light intensity dependence of CEF-driven proton pumping saturates at lower intensities than when LEF is active, possibly indicating the upper limits of the proton pumping capability of CEF, as at lower light intensities it is able to match the formation rates seen when PSII is active (**Fig. 5A**). When accounting for known and assumed H^+^/e^-^ ratios for photosynthetic electron flow, some interesting parameters can be seen. One turnover of LEF results in 3H^+^/e^-^ [65], one H^+^ from PSII, and 2 from the Q-cycle in Cyt.-b_6_f. In addition to this, a turnover of CEF through the NDH-1 complexes adds 2H^+^/e^-^, assuming a stoichiometry similar to that observed in their relatives [8] and no additional turnover of B_6_f, for a total of 5H^+^/2e^-^ from a single turnover of both LEF and CEF. With LEF inactivated by the addition of DCMU, 4H^+^/e^-^ are accounted for, with 2 from NDH-1, and 2 from Cyt.-b_6_f, resulting in 80% of the protons per turnover, with more protons moved per electron. Why is the rate of proton pumping when treated with DCMU only 40% of the no addition control (**Table 1**)? This may be able to be explained by the slower rate of complex I in other organisms (∼200 e^-^/s [63, 64])compared to that of PSII (1-400 turnovers/s [62]), along with the relative abundance of PSII compared to NDH-1 in cyanobacterial thylakoid membranes [21]. Alternatively, there may be sufficient CEF components, but limiting reductant. To test this, glucose was added to WT cells treated with DCMU, as glucose is known to increase available reductant [59], and AO quenching observed (**Fig. S4**). The addition of glucose did not increase the rate and in fact decreased it some, indicating that the limitation of CEF-driven proton pumping is not due to the lack of reductant, and is more likely due to the numbers of complexes and their respective turnover rates.

To analyze the contribution of NDH-1 complexes to proton pumping, the mutants Δ*ndhF1* and M55, lacking only NDH-1_1/2_ complexes or all NDH-1 complexes respectively, were utilized in AO assays. Interestingly, NDH-1 complexes are contributing to proton pumping even during conditions dominated by LEF since initial rates slowed to nearly 60% upon upon complete elimination of all NDH-1 complexes (in M55) and almost as much in the Δ*ndhF1* strain, though only M55 is significantly different from the WT (**Table 1**). Thus, during the dark-to-light transition, NDH-1 complexes make a strong contribution to proton pumping. In the Δ*ndhF1* strain, the remaider of the cellular complement of NDH-1 complexes is comprised of only the forms involved in CO_2_-uptake, NDH-1_3/4_. Therefore, these complexes must contribute only a minor amount to the total CEF, at least under these growth conditions. Together with the data presented here, data on the presence of the Flv1/3 system consuming electrons in their Mehler-like reaction, resulting in no additional proton pumping but acting as an electron sink to compensate for the loss of the NDH-1_1/2_ systems [66], additionally implies that the NDH-1_3/4_ complexes are incapable not only of dealing with the excess reductant but also of compensating for the loss of the NDH-1_1/2_ complexes in terms of either electron transport or proton pumping in these conditions. The proton pumping results are consistent with studies of Bernat and colleagues, showing that the respiratory forms of the NDH-1 complexes (NDH-1_1/2_) dominate the corresponding cyclic electron fluxes and that the CO_2_-uptake forms (NDH-1_3/4_) contribute less strongly [22]. The dominance and essentiality of NDH-1_1/2_ in CEF proton pumping can also be seen in **Fig. 6** and **Table 1**,where both Δ*ndhF1* and M55 are severely impaired compared to the WT when treated with DCMU. The NDH-1 null mutant, M55, was unable to form significant ΔpH in the dark-to-light transition in the time measured when treated with DCMU (**Fig. 6C**), highlighting the importance of the function of NDH-1 complexes in proton pumping under these conditions. The importance of the NDH-1_1/2_ complexes is especially highlighted by the Δ*ndhF1* mutant, which is still able to form a gradient when treated with DCMU, however at ∼5% the rate of the WT when treated with DCMU (**Fig. 6A, B**). That said, the Δ*ndhF1* strain when treated with DCMU still performs light-dependent proton pumping, indicating that deletion of only the NDH-1_1/2_ complexes is not sufficient to completely eliminate CEF-driven proton pumping, while the removal of all four NDH-1 complexes is sufficient, as seen by the absence of quenching in M55 cells treated with DCMU (**Fig. 6C**). These data indicate that the NDH-1_1/2_ complexes are the primary drivers of proton pumping in these conditions, but also imply the NDH-1_3/4_ complexes are participants as well, though to a much lesser degree than their cousins. Whether this is due to a difference in the number of NDH-1 complexes of each sort available, or some activity intrinsic to NDH-1_3/4_ complexes remains to be understood. The heavy activity of the NDH-1_1/2_ complexes in proton pumping pairs well with the established data that they are greatly active in e^-^ transport [22, 34]. Because it is known the NDH-1 complexes participate in CEF through measurements such as post-illumination fluorescence rise (PIFR) with its presence being eliminated by the deletion of NDH-1 complexes [8, 25, 67], and it is shown here that deletion of NDH-1 complexes similarly disrupts CEF-driven H^+^ pumping (**Fig. 3B, C**), indicating that the two activities are coupled by these enzyme complexes, and that NDH-1 complexes are responsible for the bulk of CEF-driven proton pumping in the dark-to-light transition. This has implications for the partitioning of electrons flowing through PSI, and it may be that the cells favor the NDH-1 complexes in times when more proton pumping, and therefore ATP production, is needed, while SDH is activated when less proton pumping is needed, but reductant must still be consumed. Given the recent structural data on the cyanobacterial NDH-1 complexes, the development of this assay system can allow for the measurement of the impact of point mutations on proton pumping, allowing for deep understandings of the mechanisms underlying their Complex I-like activities, as well as the unique CO_2_ hydration activities of the NDH-1_3/4_ complexes.

## 5. Conclusions

While it has been observed that Complex I/NDH-1 pumps protons to generate PMF in heterotrophic bacteria and archaea, mitochondria, and chloroplasts, this activity among the unique NDH-1 medley present in cyanobacteria has not been demonstrated until now. Among the processes that generate PMF, PSII and LEF are responsible for the bulk of this activity in the dark-to-light transition. The NDH-1 complexes, however, contribute to this action to a great degree, with the ability to compensate for ∼40% of the rate of proton pumping when LEF is inactivated. This is particularly important regarding the findings that fluctuating light intensity is one of the damaging stresses that photosynthetic organisms face in their natural environments [68].

## Abbreviations

AO: acridine orange
CBB: Calvin Benson Basham
CEF: cyclic electron flow
CCCP: Carbonyl cyanide *m*-chlorophenyl hydrazone
DCMU: 3-(3,4-dichlorophenyl)-1,1-dimethylurea
DCCD: *N,N*′-Dicyclohexylcarbodiimide
PAM: pulse-amplitude modulated fluorometry
PSII: photosystem II
PSI: photosystem I
COX: cytochrome oxidase
Val: valinomycin

## 6. Acknowldgements

We thank the assistance of Janet Rogers and Steven Hartson for DNA sequencing analyses within the DNA/Protein Resource Facility at Oklahoma State University. We gratefully acknowledge the financial support by the U.S. Department of Energy (DOE), Office of Science, Basic Energy Sciences, grant no. DE-FG02-08ER15968.

## Supplemental Materials

**Table S1.**
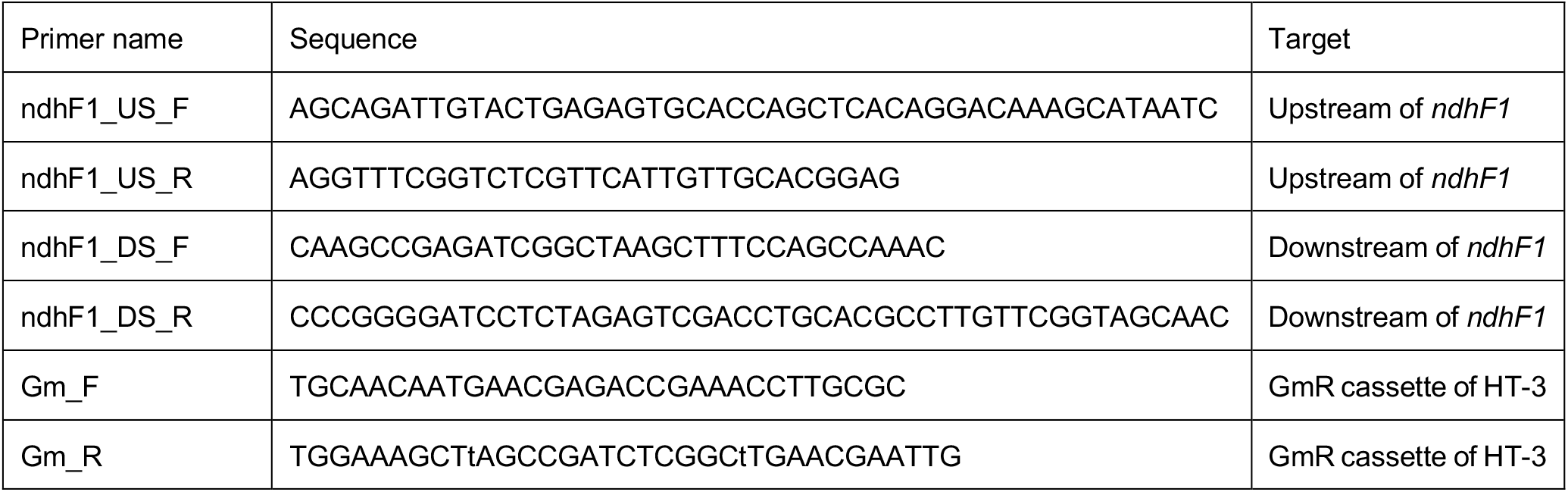
Primers used to generate Δ*ndhF1* by Gibson assembly.

**Figure S1.**
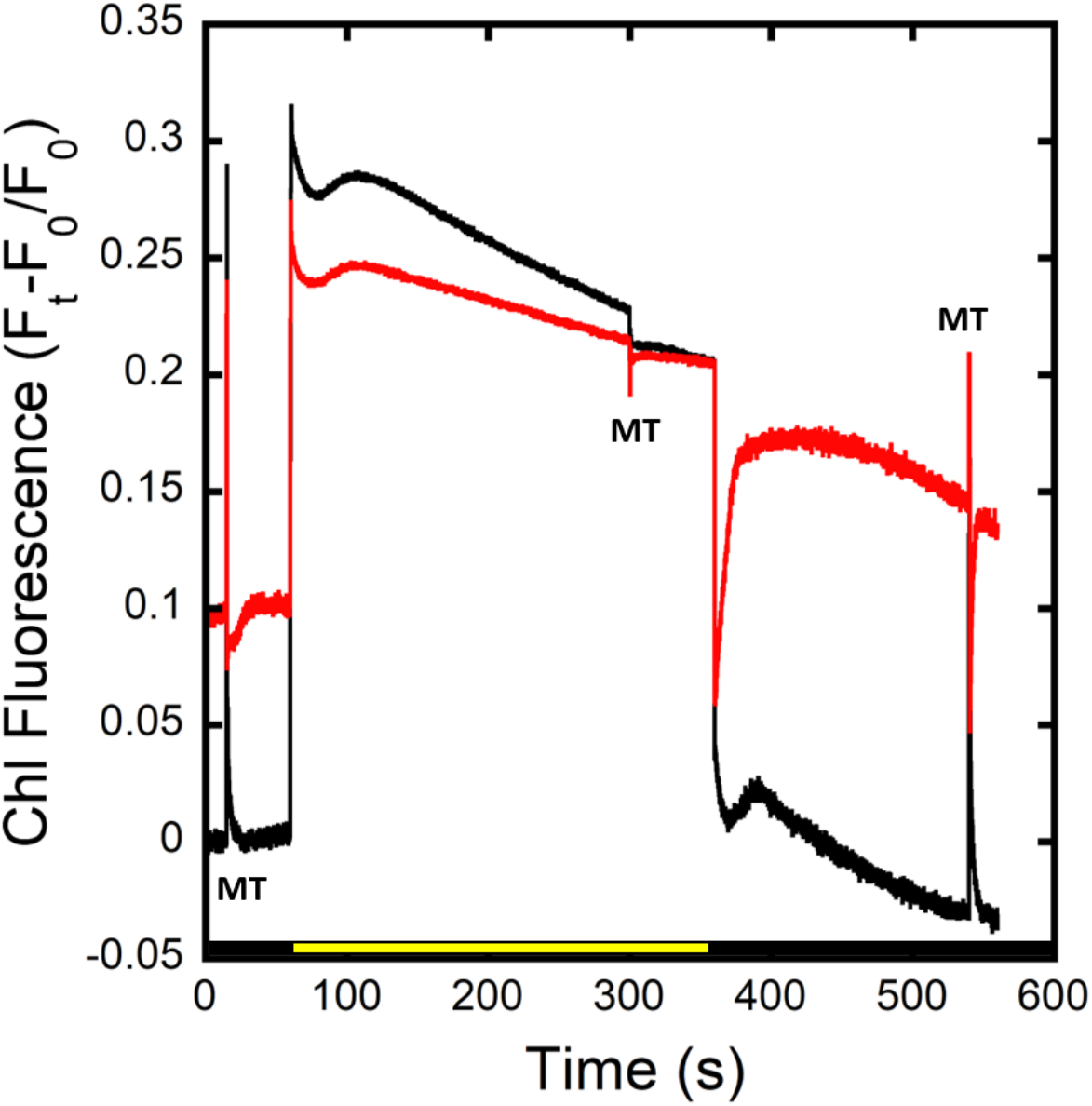
Effect of KCN on chlorophyll fluorescence in the wild type. Chlorophyll fluorescence of dark-adapted cells in TCN buffer with 150mM KCl without (black) and with (red) the addition of KCN to cells in the presence of 330μE actinic light. Multiple turnover flashes (MT) were provided where indicated. Data are normalized to F_0_ of the trace with no KCN.

**Figure S2.**
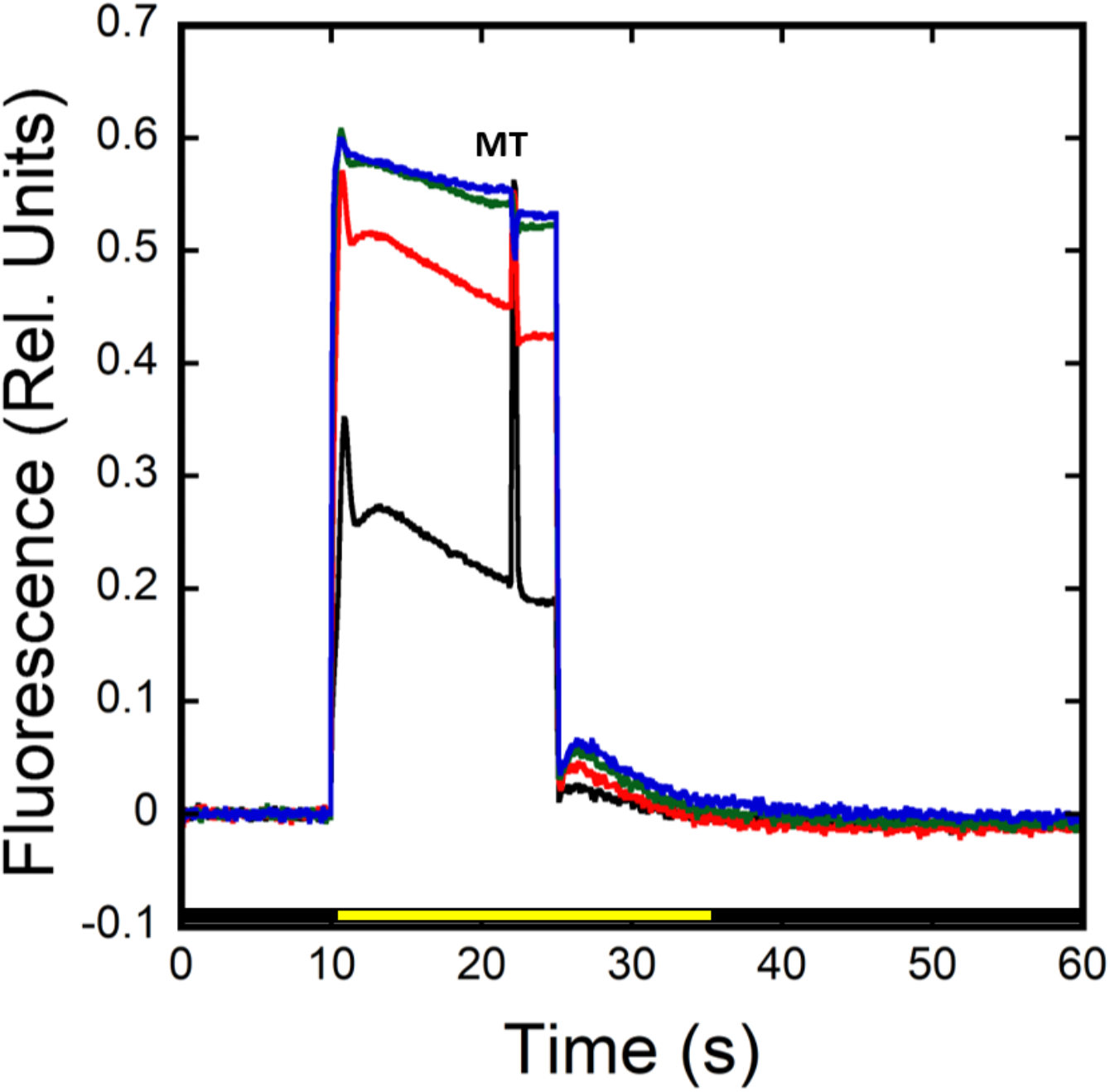
Light intensity dependence of chlorophyll fluorescence. Chlorophyll fluorescence traces were measured in dark adapted wild type cells in TCN buffer with 70uE (black), 166uE (red), 339uE (green), 660uE (blue) actinic light with a multiple turnover flash after 22 seconds of illumination. Traces normalized to F_m_ and values used to calculate q_p_ in Fig. 4B.

**Figure S3.**
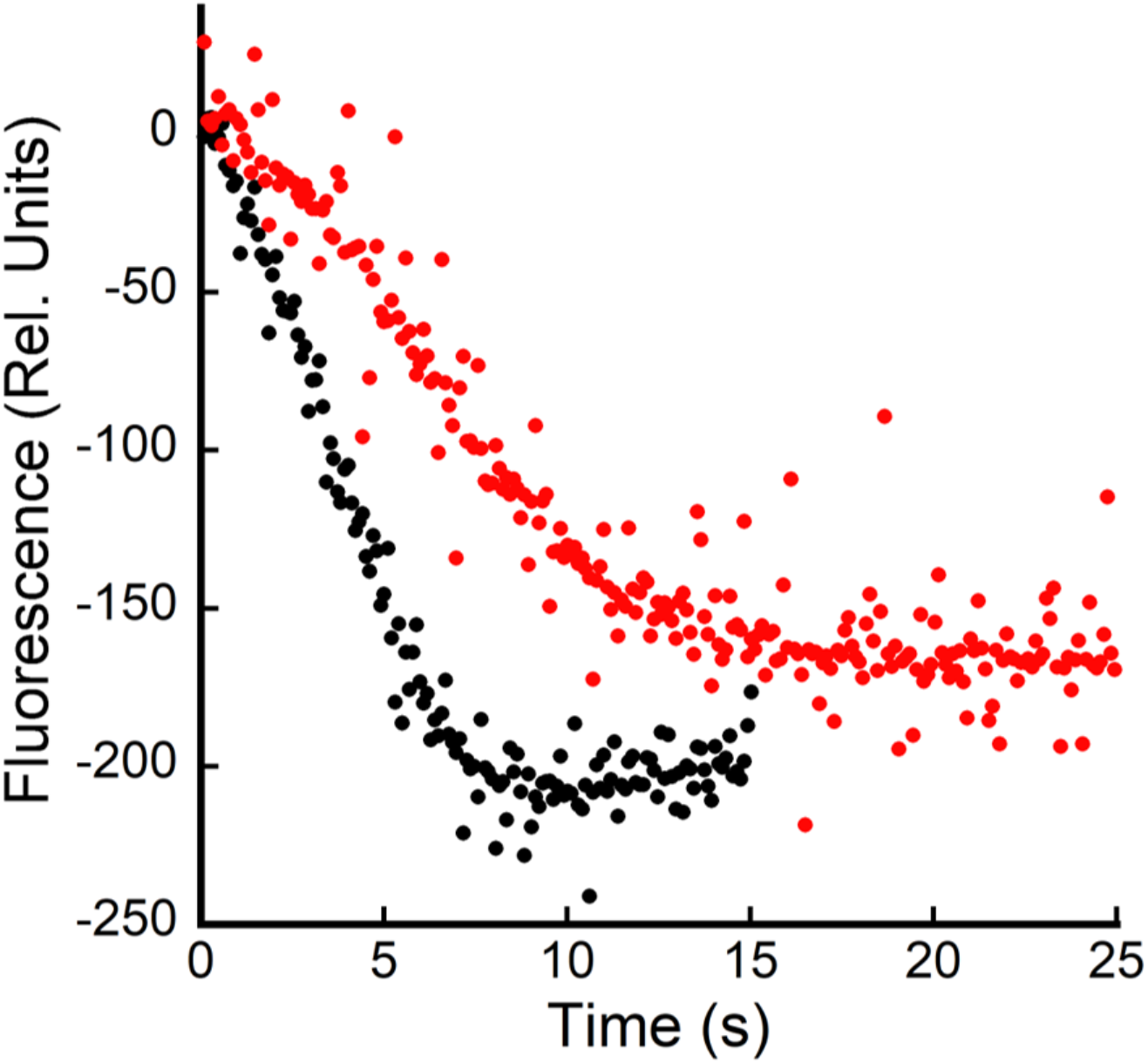
Quenching of AO fluorescence in the presence of DCMU with a longer illumination time. Acridine orange fluorescence of dark-adapted WT cells in TCN buffer with acridine orange (5µM), KCl (150mM) plus Val (10µM), DCCD (500µM), and KCN (200µM) (black) or with the addition of DCMU (red). Actinic illumination of 600µE m^−2^ sec^-1^was applied at time zero.

**Figure S4.**
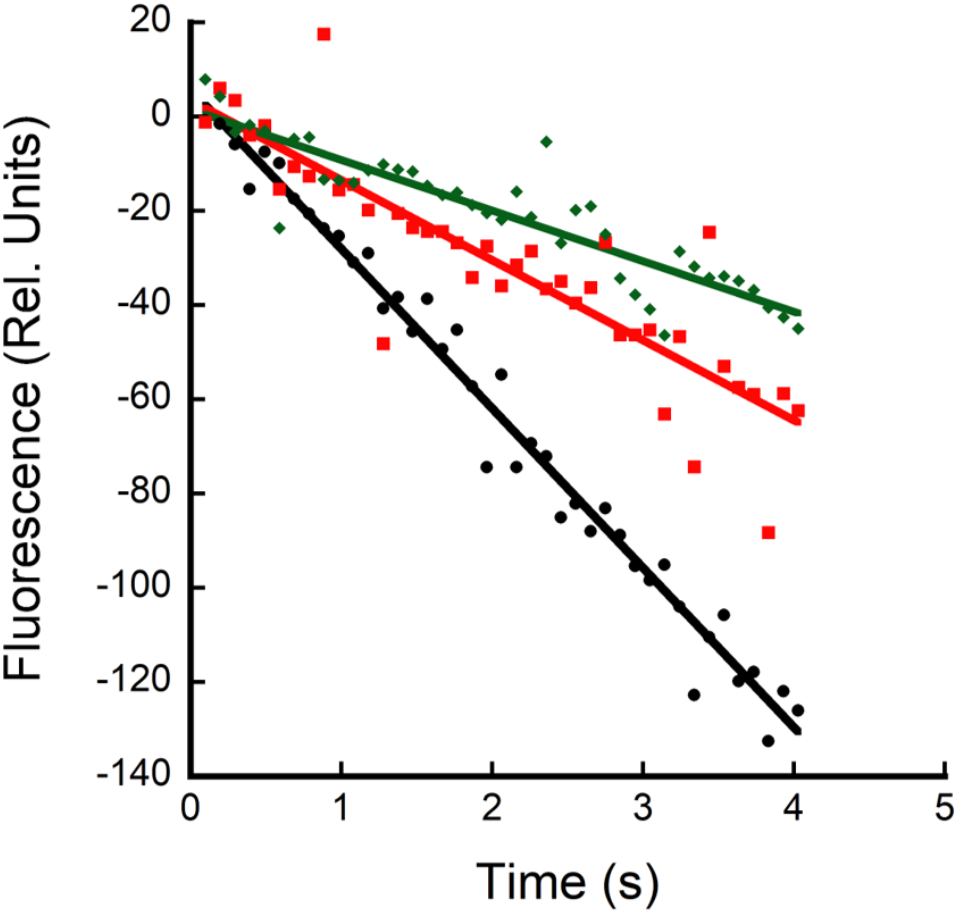
Effect of the addition of glucose on AO quenching in cells treated with DCMU. WT cells treated with background inhibitors were measured with no other additions (black circles), in the presence of DCMU (red squares), or in the presence of DCMU and glucose (green diamonds). Traces representative of three technical replicates.

**Figure S5.**
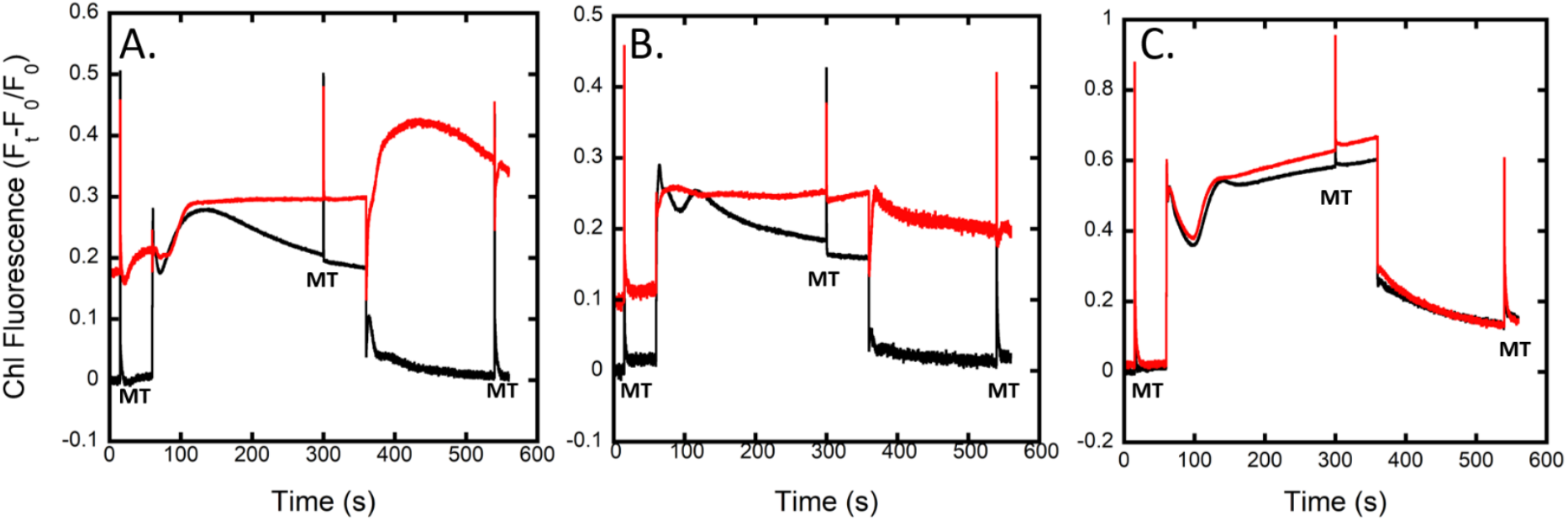
Effect of the addition of glucose on AO quenching in cells treated with DCMU. Cells of the WT (**A**), Δ*ndhF1* (**B**), and M55 (**C**) were dark adapted in TCN buffer + 150mM KCl and chlorophyll fluorescence measured with 53μE actinic illumination. Cells were either left untreated (black) or treated with 200μM KCN. Multiple turnover flashes (MT) were provided where indicated. Data normalized to F_0_ with no KCN.

